# Developmental regulators enable rapid and efficient soybean transformation and CRISPR-mediated genome editing

**DOI:** 10.1101/2025.03.03.641322

**Authors:** Anshu Alok, Vidhya Raman, Leonidas D’Agostino, Arjun Ojha Kshetry, Krishan Mohan Rai, Chunfang Wang, Samatha Gunapati, Robert M. Stupar, Gunvant B. Patil, Feng Zhang

## Abstract

Soybean transformation remains challenging and has not kept pace with the rapid advancement of genetic engineering technologies due to low efficiency, lengthy timelines, and genotype dependency. Here, we developed a streamlined transformation method by leveraging developmental regulators (DRs) to promote *de novo* shoot regeneration directly from growing soybean plants. By evaluating multiple DR combinations, our results showed that co-expression of *WUSCHEL2* (*WUS2*) and isopentenyltransferase (*IPT*) achieved higher transformation efficiencies (15.2% to 22.3%) in Williams 82 and Bert varieties than individual DRs without requiring exogenous hormones or selection agents. Moreover, this method produces heritable transgenic events within 9-11 weeks and successfully delivers CRISPR-Cas9 components, generating heritable mutations with 20% efficiency. The temporal transcriptomic and gene regulatory network analyses revealed that *WUS2*/*IPT* synergistically modulates stress responses and activates developmental pathways, orchestrating a transition from initial stress adaptation to regenerative programming. Together, our findings demonstrate that this DR-enabled approach significantly enhances soybean transformation efficiency, reduces tissue culture requirements, and offers a promising genome editing platform for soybean improvement.

## Introduction

Soybean (*Glycine max (L) Merrill*) is one of the world’s most economically important crops and an essential source of edible oil, protein, livestock feed, biofuel, and contributing to soil health through nitrogen fixation (Brar and Carter, 1993). Substantial efforts have been made over the last several decades to improve agronomically significant traits through both traditional breeding and genetic engineering approaches (Singer et al., 2023; Xu et al., 2022). In recent years, genome editing technologies, particularly CRISPR-Cas systems, have paved new ways to enable targeted genetic modifications with remarkable efficiency and precision across plant species (Gao, 2021). While various genome editing studies have shown promise in soybeans, widespread application remains limited by transformation and regeneration constraints (Freitas-Alves et al., 2024). Efficient genome editing relies on robust transformation platforms; however, recent progress in plant transformation has lagged behind the development of genome editing technologies for soybean, resulting in inefficient, genotype-dependent processes with variable levels of reproducibility (Altpeter et al., 2016).

Soybean transformation, like that of many plants, involves two critical steps: delivering DNA or RNA or protein molecules into plant cells and regenerating transformed cells into fertile plants. Molecular delivery can be achieved effectively through *Agrobacterium*-mediated transfer or direct methods like biolistic bombardment, yet plant regeneration remains the primary technical barrier (Altpeter et al., 2016). Despite progress in optimizing hormone and growth media compositions, regeneration protocols remain time-consuming and highly genotype-dependent for soybean cultivars (Xu et al., 2022).

Recent efforts to improve soybean transformation have focused on reducing tissue culture requirements and minimizing genotype dependency (Zhong et al., 2024). An emerging strategy employs developmental regulators (DRs) to enhance plant regeneration efficiency rather than relying on exogenous phytohormones (Atkins and Voytas, 2020). DR genes, typically transcription factors associated with somatic embryogenesis and de novo meristem development, can reprogram somatic cells to directly form embryos or meristematic shoots (Horstman et al., 2017). This approach was first demonstrated successfully in maize, where overexpression of *BABY BOOM* (*BBM*) and *WUSCHEL 2* (*WUS2*) induced somatic embryogenesis with dramatically enhanced transformation efficiencies across diverse genotypes (Lowe et al., 2018, 2016). Similarly, other DRs including *LEAFY COTYLEDON 1/2* (*LEC1/2*), *SHOOT MERISTEMLESS* (*STM*), and more recently, *GROWTH REGULATING FACTOR* (*GRF*) with its cofactor *GRF-INTERACTING FACTOR* (*GIF*), have shown promise in enhancing transformation efficiency and reducing tissue culture requirements across both monocot and dicot species (Debernardi et al., 2020; Kong et al., 2020; Maher et al., 2020).

The DR-based approach offers several advantages over conventional regeneration protocols: accelerated development timelines, minimal culture optimization requirements, improved transformation efficiency, reduced genotype dependency, and broader applicability across species. Furthermore, the formation of embryos or meristematic shoots serves as an intrinsic transformation marker, substantially streamlining the screening process (Atkins and Voytas, 2020). Recent studies have demonstrated the potential of DR systems in soybean, with the *GRF*-*GIF1* combination yielding up to a 2.74-fold improvement in transformation efficiency (Kong et al., 2020; Zhao et al., n.d.). However, these studies continued to rely on extended tissue culture processes using soybean primary node or half-seed cotyledonary explants as starting material. In this study, we addressed these limitations by evaluating multiple DR combinations to develop a more streamlined transformation method for soybean. Our approach aimed to establish a robust and efficient transformation protocol that eliminates both exogenous phytohormones and selection agents, thereby minimizing tissue culture requirements. Additionally, we investigated the molecular mechanisms by which DR genes reprogram cell fates through temporal transcriptomic and gene regulatory network analyses.

## Results

### Synergistic effect of WUS2 and IPT induces *de novo* shoot formation in soybean

To deliver DR genes into soybean, we built a series of T-DNA vectors, each containing three distinct functional modules: a DR expression cassette, a CRISPR-Cas9 system, and a visual marker gene cassette (Figure 1A). Within each T-DNA, we individually cloned one of three DR coding sequences: *WUS2, IPT*, or *GRF4-GIF1*. For the *WUS2* gene, we used the maize *WUS2* sequence to search against the soybean genome and identify the homologous gene, *GmWUS2* (Supplementary Figure 1A). For the *GRF4-GIF1*, we used the reported sequence from wheat to search against the soybean genome and identify the homologous gene, *GmGRF4 and GmGIF1* (Kong et al., 2020; Supplementary Figure 1B). For the *IPT*, we used the sequences reported in previous research (Maher et al., 2020). The CRISPR-Cas9 module included guide RNAs targeting the soybean major allergen gene *P34* (*Glyma.08G116300*) (Joseph et al., 2006). For non-invasive, real-time monitoring of transgenic events, we selected the *RUBY* reporter gene as a visual marker (He et al., 2020; Figure 1A). These T-DNA constructs were transformed either individually or in combinations as listed in Table 1.

**Figure 1:**
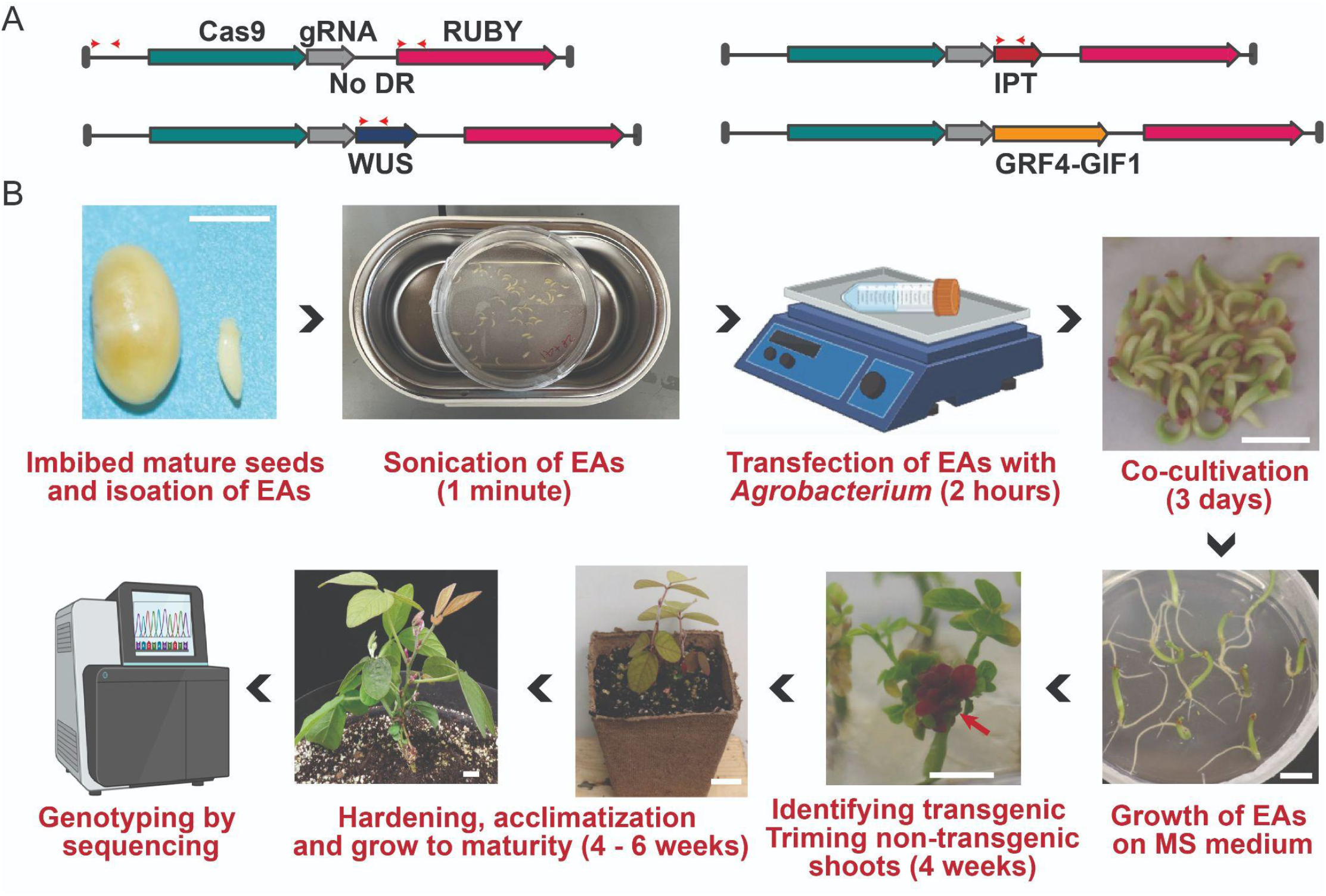
Developmental regulator (DR)-enabled soybean transformation. (A) T-DNA construct maps showing four modular components: Cas9 (green), CRISPR gRNA (gray), RUBY (ruby), and DR cassettes. The DR component includes *GmWUS2* (blue), *IPT* (red), or *GmGRF4-GIF1* (brown). Black bars indicate T-DNA borders. Red arrowheads show the locations of transgene detection primers. (B) Transformation workflow from germinating embryo isolation to transgenic/edited plant generation, with transformed plantlets displaying ruby-colored shoots (red arrow). Scale bar: 1 cm.

**Table 1:** DR-mediated soybean transformation efficiencies.

| DRs | Soybean variety | # of explants | # of explants with transgenic shoots | Transformation efficiency (%) | Average transformation efficiency (%) |
| --- | --- | --- | --- | --- | --- |
| No DRs | W82 | 27 | 0 | 0 | 0 |
| No DRs | W82 | 20 | 0 | 0 |  |
| No DRs | W82 | 10 | 0 | 0 |  |
| IPT | W82 | 20 | 0 | 0 | 6.4 |
| IPT | W82 | 35 | 5 | 14.3 |  |
| IPT | W82 | 40 | 2 | 5 |  |
| WUS | W82 | 20 | 1 | 5 | 3.3 |
| WUS | W82 | 20 | 0 | 0 |  |
| WUS | W82 | 40 | 2 | 5 |  |
| WUS/IPT | W82 | 26 | 8 | 30.8 | 22.3 |
| WUS/IPT | W82 | 74 | 12 | 16.2 |  |
| WUS/IPT | W82 | 25 | 5 | 20 |  |
| WUS/IPT | Bert | 35 | 4 | 11.4 | 15.2 |
| WUS/IPT | Bert | 37 | 7 | 18.9 |  |
| GRF | W82 | 30 | 0 | 0 | 0 |
| GRF | W82 | 12 | 0 | 0 |  |
| WUS/GRF | W82 | 24 | 2 | 8.3 | 5.8 |
| WUS/GRF | W82 | 30 | 1 | 3.3 |  |

While various explant types have been used for soybean transformation, including cotyledonary nodes from immature and mature seeds, half-seeds, hypocotyls, primary nodes, immature embryos, and mature embryonic axis(Xu et al., 2022), we chose to use 2-day germinating embryos (i.e. embryonic axis or EA) based on two key considerations: 1) during primary shoot development, the DRs could promote *de novo* axillary shoot formation on the explants(Maher et al., 2020), and 2) the transformed embryos could continue to grow into plantlets without exogenous growth hormones. Our transformation protocol involved isolating germinating embryos from imbibed mature seeds, followed by transformation using *Agrobacterium* strain AGL1 carrying the T-DNA constructs (Figure 1B). After a three-day co-cultivation period, the treated embryos were cultured on Growth medium I and II (Supplementary Table S1) without growth hormones or selection reagents. The transformed embryos germinated and produced multiple shoots, averaging 6-8 shoots per explant. A subset of these explants exhibited 1-2 emerging *RUBY*-colored shoots (Figure 1B). The plantlets with *RUBY*-colored shoots were then transplanted to soil and grown as T0 plants to maturity, resulting in T1 seed production (Figure 1B). These transgenic shoots were readily identified by their *RUBY* coloration throughout development.

We evaluated the transgenic shoot formation efficiency with DR genes in the soybean variety *Williams 82*. The efficiency was calculated as the percentage of explants developing *RUBY*-colored shoots relative to the total number of treated explants. In single-DR treatment groups, transformation efficiencies were 3.3% with *WUS2*, 6.4% with *IPT*, and 0% with *GRF4-GIF*, demonstrating that individual DRs—particularly *IPT*—could induce *de novo* shoot formation (Table 1). Notably, the dual-DR combination of *WUS2* and *IPT* exhibited substantially higher transformation efficiency, averaging 22.3%, while the *WUS2* and *GRF4-GIF1* combination showed 5.8% efficiency. These results indicate that DR combinations including *WUS2* significantly enhanced transgenic shoot formation, with the *WUS2*/*IPT* combination substantially outperforming all other DR groups (Table 1). To validate this finding across genotypes, we applied the *WUS2*/*IPT* combination to another soybean variety, *Bert*, and observed a comparable transformation efficiency of 15.2%.

Among these transgenic shoots, we observed varying intensities of *RUBY* coloration in leaves, flower petals, seed pods, and seed coats (Figure 2A). To confirm that the *RUBY*-colored shoots were genuine transformation events, we sampled nine T0 plants and verified the presence of transgenes using polymerase chain reaction (PCR) (Figure 2B). Additionally, using DR gene-specific primers, we found that all nine T0 plants contained *WUS2* and *IPT* transgene sequences (Supplementary Figure 2A). All *RUBY*-positive shoots from these T0 plants produced seeds. However, the *RUBY*-colored seed pods and seeds were noticeably smaller compared to those from non-transgenic sibling shoots (Figure 2A). To assess transgene inheritance, we grew 10 T1 seeds collected from two independent *RUBY*-positive T0 plants and performed PCR-based genotyping. Our results revealed differential transgene transmission rates, with 8 out of 10 T1 plants from one T0 parent and 3 out of 10 T1 plants from the other showing successful transgene inheritance (Figure 2B, Supplementary Figure 2B).

**Figure 2:**
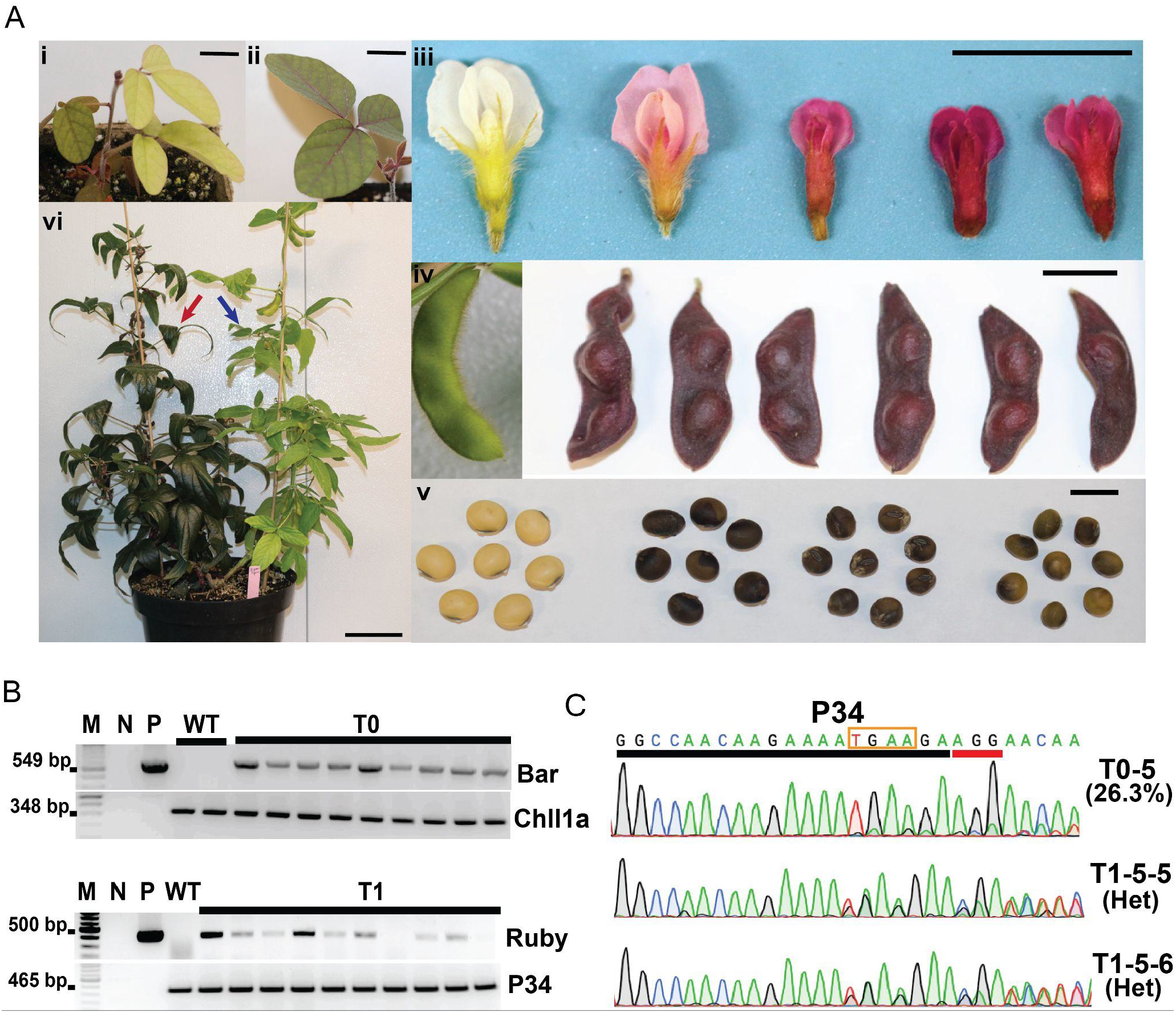
Characterization of transgenic and gene-edited soybean plants. (A) Phenotypic analysis of T0 plants showing transgenic versus non-transgenic tissues: independent transgenic shoots with varied ruby coloration (i-ii), flowers (iii), seed pods (iv), and seeds (v). A chimeric plant displaying both transgenic (red arrow) and non-transgenic shoots (blue arrow) (vi). Scale bar: 1 cm (i - v); 10 cm (vi). (B) PCR genotyping of T0 and T1 plants. Transgene integration confirmed in nine T0 plants using the T-DNA primers (Bar) with the *ChlI1a* gene primers as internal control, and in ten T1 progenies using the RUBY primers with the *P34* gene primers as internal control. M: 1 kb plus DNA ladder; N: no-DNA control; P: plasmid control; WT: non-transgenic wild type control; T0/T1: individual plant samples. (C) Sanger sequencing of the *P34* target site in the T0 plant, T0-5. Two T0-5 progenies (T0-5-5 and T0-5-6) exhibited heterozygous mutations with double peaks in sequencing chromatograms. The P34 target sequence on top shows 20-bp CRISPR site (black bar), 3-bp PAM (red bar) and 4-bp deletion region (brown box) found in T1 mutants.

### Heritable CRISPR-Cas9 mutations generated from DR-induced shoots

In addition to evaluating DR gene inheritance, we assessed the efficiency of CRISPR-mediated genome editing, as each DR-expressing T-DNA construct contains a CRISPR-Cas9 cassette targeting the *P34* allergen gene. The target regions were PCR amplified from nine individual T0 plants and analyzed using the Next Generation Sequencing. Sequence analysis revealed that 6 of 9 T0 plants carried editing events with efficiencies ranging from 0.1% to 34.5% (Supplementary Figure 3A). To examine mutation heritability, we screened 10 T1 plants derived from event T0-5, which showed a 26.1% mutagenesis rate, using PCR and Sanger sequencing (Figure 2C). Among these T1 plants, two heterozygous mutants were identified with 4-bp deletion at the target site (Figure 2C, Supplementary Figure 3B). These results demonstrate that our transformation method effectively delivers CRISPR-Cas9 and induces heritable mutations in soybean. Together, our findings establish that combining *WUS* and *IPT* dramatically enhances soybean transformation efficiency through promotion of de novo shoot formation. This hormone-free approach achieves the transformation timeline to 9-11 weeks from initial embryo isolation to T1 seed production. By simplifying the transformation process while maintaining high editing efficiency, this method promises to accelerate soybean genetic engineering efforts.

### Transcriptional dynamics of *WUS2*/*IPT* synergy during de novo shoot formation

The synergistic effect of *WUS2* and *IPT* on *de novo* shoot formation efficiency prompted us to investigate the transcriptional dynamics induced by these DR genes. We performed bulk RNA sequencing at 0, 3, and 6 days after transformation (DAT) using four treatments: empty no-DR vector control (EV), *WUS2* alone, *IPT* alone, and *WUS2*/*IPT* combination. Principal component analysis (PCA) revealed distinct temporal separation of transcriptional profiles at 3 and 6 DAT (Supplementary Figure 4A). At 3 DAT, samples clustered primarily by treatment with high concordance between replicates (Supplementary Figure 4B). By 6 DAT, samples were segregated into two distinct clusters: one comprising EV and *WUS2* treatments, and another containing *IPT* and *WUS2*/*IPT* treatments (Supplementary Figure 4C). This clustering pattern was further supported by distance matrix analysis (Supplementary Figure 4D). To delve into these expression patterns, we performed K-means clustering of differentially expressed genes (DEGs), which revealed five distinct clusters at 3 DAT and two at 6 DAT (Figure 3A and B; Source data file). At 3 DAT, we identified synergistic expression of genes regulating distinct yet interconnected pathways associated with early DR response. Gene Ontology (GO) enrichment analysis revealed that genes involved in cell cycle progression, hormone response, cell wall remodeling, and meristematic tissue establishment were highly upregulated in *WUS2*/*IPT* treatment (Clusters 3 and 4; Figure 3A; Source data file), suggesting initiation of somatic cell reprogramming toward a regenerative state. Specifically, genes involved in DNA replication and cytokinin pathway activation were significantly expressed in *WUS2*/*IPT*. In contrast, stress response, defense, and light-responsive genes were highly upregulated in EV, *IPT*, and *WUS2* treatments (Clusters 1 and 2; Figure 3A; Source data file). Notably, coordinated expression of *WUS2*/*IPT* not only suppressed stress and defense response genes but promoted cell cycle activation and hormone response (Supplementary Figure 5A).

**Figure 3:**
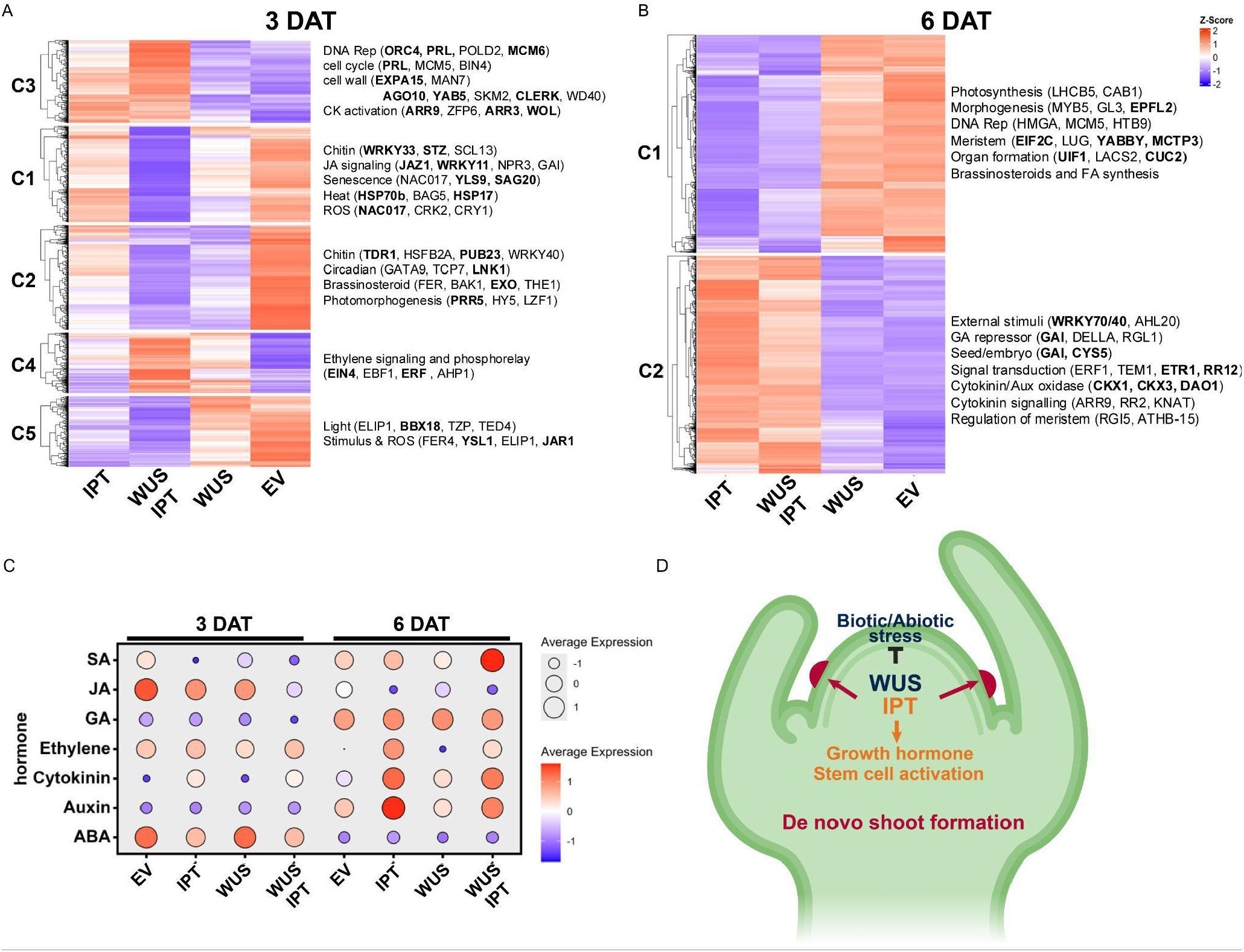
Transcriptional dynamics of DR-transformed soybean shoots. K-means clustering analysis heat maps at 3 DAT (A) and 6 DAT (B), with enriched GO terms and genes highlighted per cluster. (C) Gene pathway enrichment analysis of differentially expressed genes involved in plant hormone pathways. (D) Schematic model showing the synergistic effects of *WUS2* and *IPT* on *de novo* shoot formation through the inhibition of biotic/abiotic stress responses and promotion of growth hormone and stem cell activation pathways.

The transcriptional landscape evolved further at 6 DAT, where two major clusters emerged, with *IPT* and *WUS2*/*IPT* showing similar DEG patterns distinct from *WUS2* and EV (Figure 3B; Source data file). This is consistent with the observation that *IPT* played a dominant role in activating genes involved in cytokinin signaling, meristem regulation(Nguyen et al., 2021). Key genes involved in stem cell activation, including *RGI5, ATHB-15*, and cytokinin targets such as *ARR9, ARR2*, and *KNAT*, were upregulated in *WUS2*/*IPT*(Baima et al., 2001; Dai et al., 2017; Fernandez et al., 2020). Additionally, negative regulators of GA response, including DELLA subfamily GRAS regulatory genes (*GAI, RGL1*), were upregulated in both *WUS2*/*IPT* and *IPT* treatments. Interestingly, brassinosteroid-associated genes (*FER, BAK1, EXO, THE1*) were downregulated in *WUS2*/*IPT* compared to EV at both timepoints (Figure 3B; Source data file)(Hamant et al., 2002).

To further understand the role of hormone signaling in this process, we conducted a targeted principal component analysis of hormone-responsive genes (Figure 3C and Supplementary Figure 5B; Source data file). This analysis revealed a clear temporal transition in hormone profiles. At 3 DAT, samples clustered closely and showed strong associations with stress-related hormones, including jasmonic acid (JA), abscisic acid (ABA), and ethylene, suggesting an initial biotic stress response to *Agrobacterium* co-cultivation (Figure 3C and Supplementary Figure 5B; Source data file). By 6 DAT, a marked shift occurred: *WUS2*/*IPT* and *IPT* treatments showed stronger induction of auxin, and cytokinin pathways that are known to be associated with *de novo* shoot development (Figure 3C and Supplementary Figure 5B; Source data file). This temporal shift from stress response to regenerative processes highlights the dynamic role of DRs in orchestrating both initial stress adaptation and subsequent developmental reprogramming. Notably, the distinct hormone profiles between *WUS2*/*IPT* and other treatments at 6 DAT align with their different regeneration efficiencies, highlighting the importance of hormone balance in shoot formation. Together, our transcriptional analysis reveals that *WUS2*/*IPT* expression initiates a complex developmental program: it initially prepares explants for tissue dedifferentiation by modulating stress responses and cell fate genes, and then promotes *de novo* shoot formation through coordinated regulation of meristem activity and growth hormone signaling networks (Figure 3D).

## Discussion

In this study, we developed a rapid and efficient soybean transformation method by DR-induced *de novo* shoot formation. Our approach demonstrates several key improvements compared to conventional transformation methods. First, we fully eliminated the use of exogenous hormones and selection agents, whereas traditional methods using half seed or embryonic axis require these components, thereby simplifying tissue culture requirements (Cho et al., 2022; Zhao et al., n.d.; Zhong et al., 2024). Second, this DR-enabled approach eliminates the need for separate shooting and rooting processes. The rooting step can be particularly challenging for different soybean cultivars and often extends the process by more than 4 weeks. By bypassing this requirement, our method is substantially shorter than widely-used soybean transformation methods (B et al., 2020; Cho et al., 2022). Third, the combination of *WUS2* and *IPT* achieved significantly higher transformation efficiency (22.3%) compared to single DRs or other combinations, suggesting a synergistic effect in promoting shoot regeneration. Additionally, this method enabled efficient CRISPR-mediated genome editing with frequencies up to 33.0% in T0 plants and heritable mutations in 20% of T1 plants. Collectively, these improvements, along with successful application in two soybean varieties, represent a significant advancement that could accelerate soybean genetic engineering for both research and agricultural applications.

Despite these promising results, several aspects of this method require further optimization. The strong expression of the *RUBY* reporter gene appears to negatively impact plant growth, potentially affecting gamete viability and seed development, as evidenced by reduced seed size in transgenic lines. Alternative reporter genes, such as fluorescent proteins, could be evaluated for tracking transgenic shoots (Baysal et al., 2025). The variable transgene transmission rates among T0 plants also suggest potential chimeric tissue formation. Furthermore, the constitutive overexpression of *WUS2* and *IPT* may adversely affect plant development, suggesting the need for strategies to precisely regulate these genes, such as using inducible promoters or implementing site-specific recombination systems to remove them post-transformation. In addition, integration with newly developed genome editing systems, such as base editing and prime editing, could enable more precise genome modifications. These refinements will be imperative for further enhancing the utility of this method in soybean genetic engineering applications.

To understand the molecular basis of the synergistic effect of *WUS2*/*IPT* on enhanced shoot formation efficiency, we performed temporal transcriptome analysis on samples with *WUS2* and *IPT* individually or in combination. Notably, genes associated with stress response were upregulated in the no DR genes control group, while these genes were significantly downregulated with the synergistic activity of *WUS2/IPT*. The early suppression of stress-related genes is intriguing, as stress responses due to *Agrobacterium* infection often hinder transformation efficiency. This finding suggests that *WUS2*/*IPT* not only promotes shoot regeneration but also mitigates transformation-associated stress, potentially contributing to the higher success rate observed in our experiments.

Additionally, genes involved in cell cycle progression and hormone regulation, followed by shoot meristem development, were differentially expressed in *WUS2/IPT* and *IPT* treatments. The hormone-responsive gene analysis further illuminated the temporal dynamics of cellular reprogramming. The transition from stress-related hormones (JA, ABA, ethylene) at 3 DAT to growth-promoting hormones (auxin, cytokinin) at 6 DAT indicates a well-coordinated developmental program. Importantly, the distinct hormone profiles between *WUS2*/*IPT* and other treatments at 6 DAT align with their different regeneration efficiencies, highlighting the importance of hormone balance in shoot formation. Together, our study provides a practical solution for soybean transformation while advancing the basic understanding of plant regeneration. The success of this DR-based approach suggests similar strategies might be applicable to other recalcitrant soybean genotypes or grain legume species. Future work will focus on implementing alternative reporter genes and optimizing DR expression to improve transformation efficiency while minimizing developmental impacts.

## Materials and Methods

### Construction of T-DNA vectors

All the vectors were cloned using the Golden Gate assembly approach, which was based on modules A, B, C’ and D (Čermák et al., 2017). Module A carries the Cas9 coding sequence driven by the *GmUbi* promoter, whereas Module B contains CRISPR guide RNA spaced driven by the *CmYLCV* promoter. The coding sequences of each developmental regulator gene, *GmWUSCHEL*2 (*WUS2*), Isopentenyltransferase (*IPT*), and *GmGRF4-GIF1* were cloned into module C’ (Supplementary Figure 6). A *RUBY* visual reporter gene was cloned under *At*Ubi10 constitutive promoter in module D. Further, these modules were assembled into a destination vector, pTRANS_231d and pTRANS_230, following the Golden Gate assembly method (Čermák et al., 2017).

### Seed sterilization and explant preparation

Mature dry seeds of soybean (cv. *Williams 82* and *Bert*) were surface sterilized for 16 h using chlorine gas, produced by mixing 5 ml of Hydrochloric Acid (HCl, TraceMetal™ Grade, Thermo Fisher) with 100 mL of commercial bleach. Sterilized seeds were placed on germination medium plates (Supplementary Table S1) in the dark at room temperature for 6-7 hours. Further, seeds were taken out from the germination plate and soaked in sterile water overnight at room temperature. Intact embryonic axes were isolated and undeveloped leaves were broken carefully without damaging the meristematic region. All isolated embryonic axes were kept in Co-cultivation medium (Supplementary Table S1) before *Agrobacterium* transformation.

### Preparation of *Agrobacterium* suspension

The T-DNA vectors were transformed into the AGL1 strain of *A. tumefaciens* using the freeze-thaw method and plated onto YEP agar containing rifampicin 10 μg/mL and kanamycin 50 μg/mL. A single colony was inoculated in 20 mL of YEP liquid medium (rifampicin 10 μg/mL and kanamycin 50 μg/mL) and cultured at 28°C overnight with shaking at 220 rpm. The bacterial culture was centrifuged at 4000 rpm, for 10 min at 21°C, and the pellet was resuspended in the co-cultivation medium (Supplementary Table S1). Further, *Agrobacterium* suspension was incubated at room temperature in an orbital shaker at 60 rpm for about 2 hours in the dark, and then OD600 of the cultures was adjusted to 0.5 before infection. For co-transformation with two constructs, equal volumes with equal OD600 of each culture were mixed before transformation.

### *Agrobacterium* mediated transformation of germinating embryo

Isolated EAs were mixed with 15ml *Agrobacterium* suspension in a petri dish and sealed with parafilm. The sealed plate was subjected to sonication in water (Sonicator, Fisher Scientific Model-FS6) for 1 min. After sonication, another 15 ml of *Agrobacterium* suspension was added to each plate in a laminar flow hood. The plate was covered with foil and kept on a shaker at 60 rpm for 2 hours at room temperature. After inoculation, excess bacterial suspension was removed and embryonic axis (EAs) were transferred to a single layer of autoclaved sterile filter paper in a 15 × 100 mm petri dish. The filter paper was moistened with 700µl of co-cultivation medium (without *Agrobacterium*). Plates were sealed with parafilm and kept in a 22°C transparent incubator for 3 days under low light (10 μmol m^−2^ s^−1^). After 3-days of co-cultivation, EAs were rinsed with co-cultivation medium with cefotaxime (200 μg/mL) and timentin (100 μg/mL), and the base of each embryonic axis was embedded in Growth medium I (Supplementary Table S1) at 24 °C with a 16/8-hour light/dark cycle. After two weeks, the apical elongated stem was trimmed from each explant and transferred to Growth medium II (Supplementary Table S1) for the emergence of transformed pink shoots.

### Hardening acclimatization and plant growth condition

Soybean plantlets with transgenic shoots were removed from plant tissue culture containers and gently washed to remove agar gel without damaging the roots. Plants were transplanted to 4-inch biodegradable peat pots containing potting mix soil and covered with transparent plastic covers to maintain humidity. After 10-14 days, the covers were gradually removed. After 15 days, the 4-inch pots were transferred into 5-gallon pots. The plants were grown in growth chamber with temperature maintained at 26 °C, 55% relative humidity, and a 16/8-hour light/dark cycle until maturity. Plants were continuously watered and fertilized at one-week intervals.

### DNA extraction, PCR genotyping, and gene editing mutation detection

DNA was extracted from the T0 and T1 plants using the bead grinder and CTAB method (Allen et al., 2006). Polymerase chain reaction (PCR) was performed from the genomic DNA with the oligonucleotides listed in Supplementary Table S2 by using Q5 high fidelity DNA Polymerase as per manufacturer protocol (New England Biolab, MA). For T0 plants, PCR amplicons of the *P34* gene target site were sequenced using Illumina paired-end read sequencing (Genewiz Inc., South Plainfield, New Jersey, USA). Sequencing reads and mutation rates were analyzed by CRISPResso2 using the default setting for CRISPR-Cas9 analysis(Clement et al., 2019). To detect mutations in T1 plants, the PCR product from the *P34* gene of each plant was subjected to Sanger sequencing. The samples showing double peaks near the target site were further confirmed by Nanopore-sequencing (Plasmidsaurus, OR) and analyzed by CRISPResso2(Clement et al., 2019).

### Sample collection and RNA-seq

Explants transformed with vectors having *WUS2, IPT, WUS2/IPT*, and Empty Vector (EV) were subjected to sampling. Infection, co-cultivation and transfer of embryonic axis were carried out as described above. At zero days, three days or six days after transformation, shoot regions (4-6 mm) were excised from corresponding explants and immediately frozen in liquid nitrogen for RNA extraction. For each time point, five excised shoot samples were pooled for one biological replicate, and three biological replicates were collected for each treatment. Total RNA was extracted using a RNeasy Plant Mini Kit (Qiagen, CA) according to the manufacturer’s protocol, including an on-column DNase digestion. Library preparation and Illumina sequencing were carried out at Novogene using the NovaSeq 6000 PE150 (Novogene, Sacramento, CA).

### RNAseq data analysis

Raw reads were processed using fastp (0.23.2) with the “-detect_adapter_for_pe”, “-trim_poly_x”, and “-trim_poly_g” parameters to remove adapters and filter bad reads. Filtered reads were aligned to the *Glycine max* reference genome (Wm82.a4.v1) downloaded from https://phytozome-next.jgi.doe.gov using kallisto (0.48.0). Kallisto abundance files were used to perform differential analysis in R using DESeq2 (1.44.0) with the threshold of p.adj < 0.05. Variance stabilized transformation (VST) values for gene expression were used for sample-wise PCA, distance matrix plotting, and hormonal PCA. Z-scaled VST expression values were used for k-means clustering and visualization and venn diagrams. Heatmaps were generated using the “pheatmap” function in ComplexHeatmap (2.20.0). Hormonal PCA plot was generated using the “fviz_pca_biplot” function in factoextra (1.0.7).

## Data availability

The gene IDs or accession numbers include *AtWUS2* (AT2G17950.1), *GmWUS2* (Glyma.01G166800.1), *AtGRF4* (AT3G52910), *GmGRF4* (Glyma.12G014700.1), *TaGRF4* (TraesCS6A01G269600) and the soybean *P34* gene (Glyma.08G116300). The sequence information of the DRs is available in Supplementary Figure 6. The sequence information of the oligos is available in Supplementary Table S2. The Next-Generation sequencing data generated in this study have been deposited in the NCBI Sequence Read Archive (SRA) under accession code PRJNA1219159. The RNA-seq data have been deposited under accession code PRJNA1218798. The processed CRISPResso2 output files for each sample are available from github [https://github.com/ZhangLab-UMN/SoybeanDR_NGS]. Other data generated in this study are provided in the Supplementary Information and Source Data file.

## Acknowledgments

A.A., V.R., C.W., and F.Z. are supported by the National Science Foundation (IOS-2040218 and IOS-2206920) awards. F.Z. were supported by USDA NIFA award #2021-67013-34565. L.D., A.O.K., and G.P. are supported by the Texas Governor’s University Research (GURI) grant. We thank all the Zhang lab members for their inputs. We thank Dr. Danial Voytas and Dr. Colby Starker, University of Minnesota, for sharing the modular cloning kit.

