## Supplementary information for "Developmental regulators enable rapid and efficient soybean transformation and CRISPR-mediated genome editing"

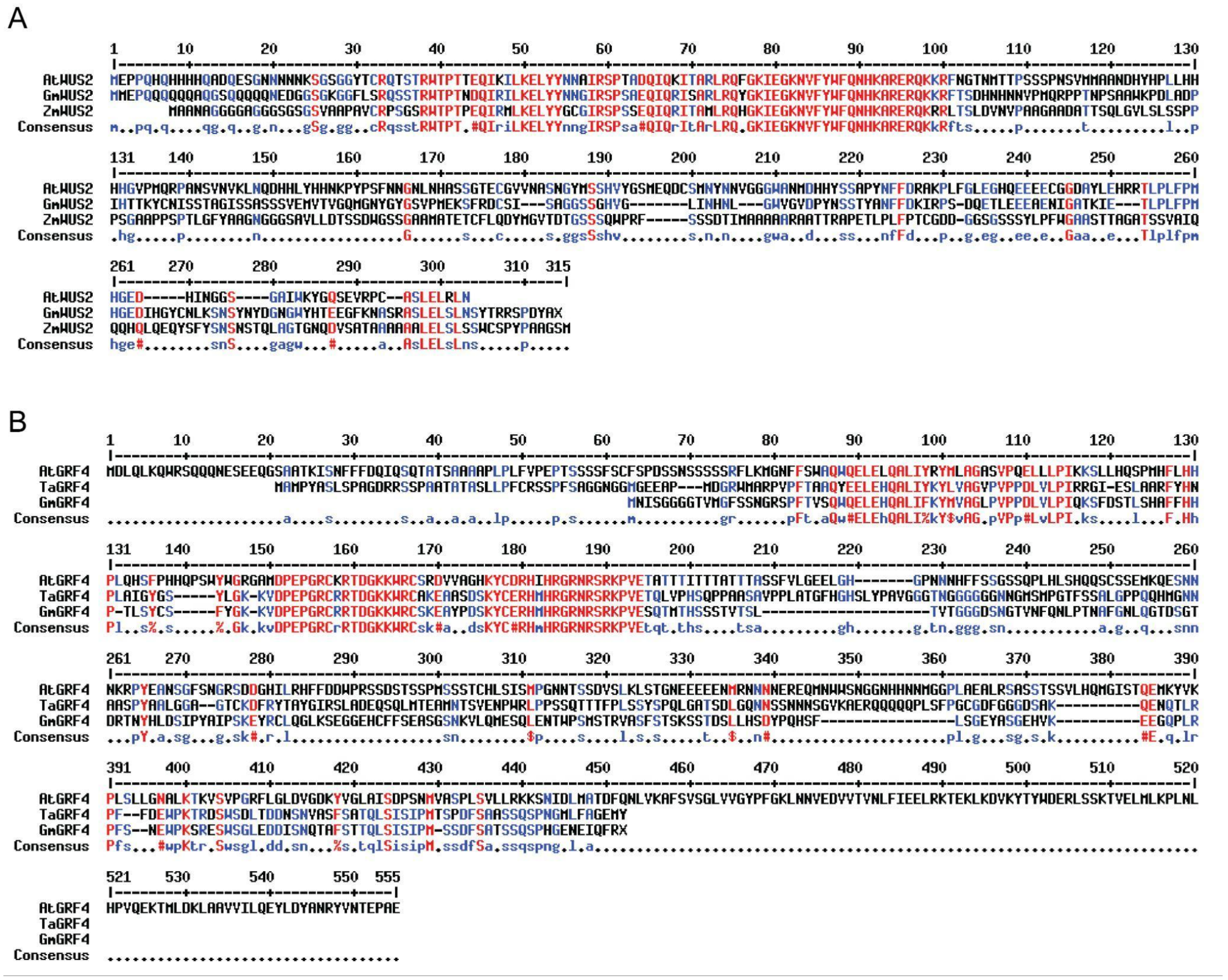


**Supplementary Figure 1:** Multiple sequence alignment of WUS2 and GRF4 protein from *Arabidopsis thaliana*, *Zea mays, Triticum aestivum* and *Glycine max*.

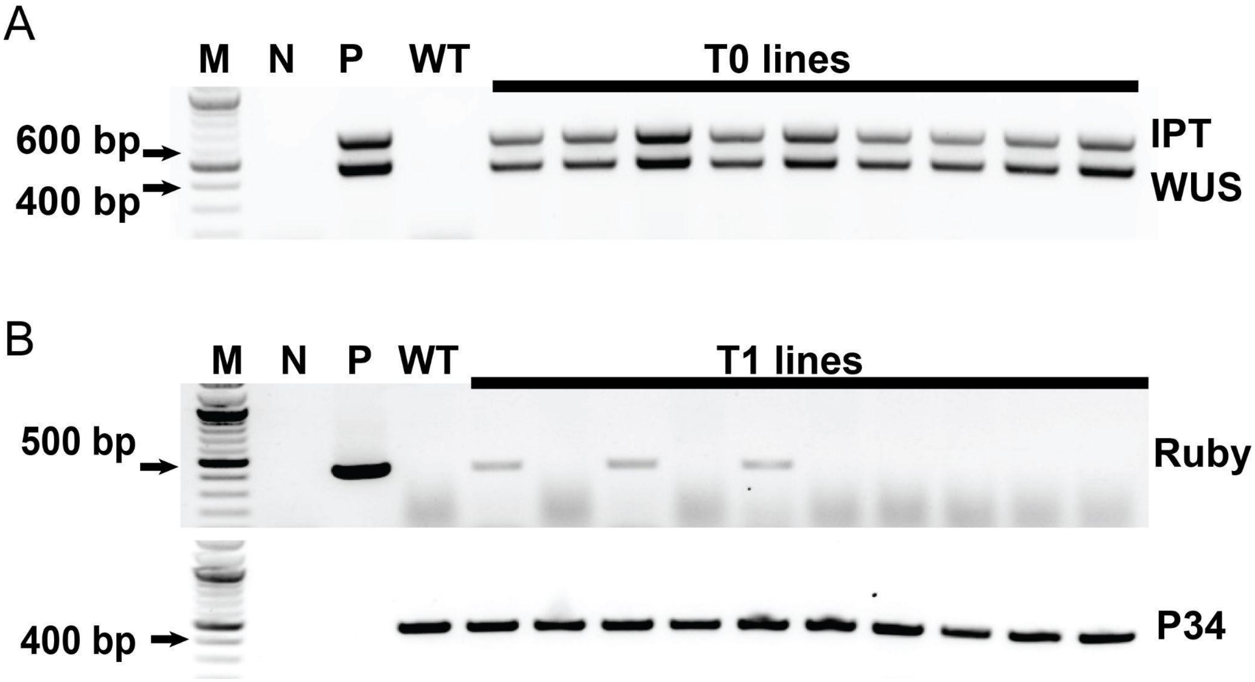


**Supplementary Figure 2: Characterization of transgenic soybean plants.** (A) PCR genotyping of T0 plants. Transgene integration of DRs confirmed in nine T0 plants using the WUS and IPT primers, and (B) T-DNA segregation in ten T1 progenies using the RUBY primers with the P34 gene primers as internal control. M: 1 kb plus DNA ladder; N: no-DNA control; P: plasmid control; WT: non-transgenic wild type control; T0/T1: individual plant samples.

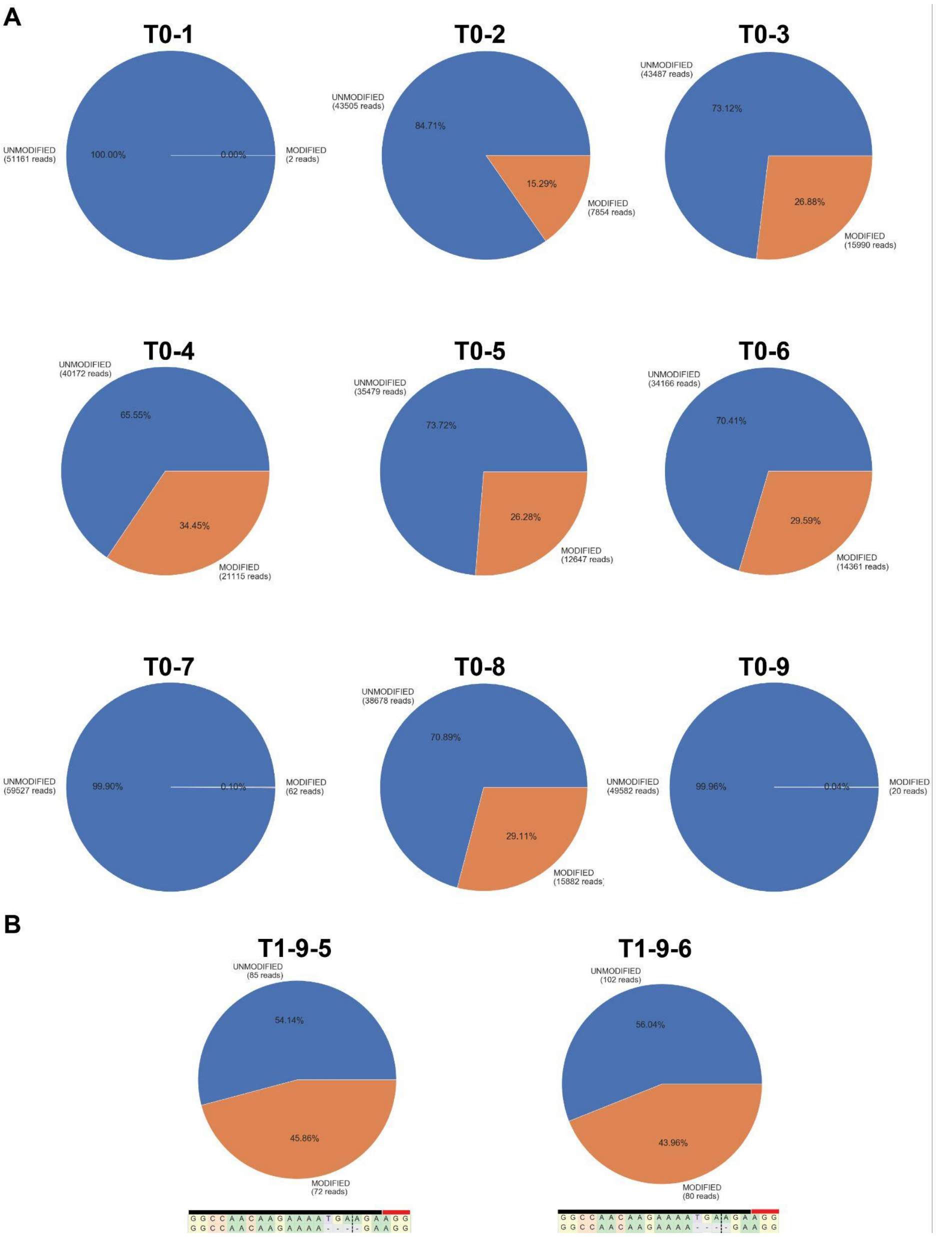


**Supplementary Figure 3:** Characterization of CRISPR-mediated gene editing in T0 and T1 plants. Next-generation sequencing analysis of mutation rates at the P34 gene in nine T0 plants (A) and two T1 plants (B). Pie charts generated by CRISPResso2 show the proportion of wild-type (blue) and edited (orange) sequences for each plant. (B) Sequence alignment of heterozygous mutations, with the target site (black bar), PAM sequence (red bar), and 4-bp deletion region (dashed lines) indicated.

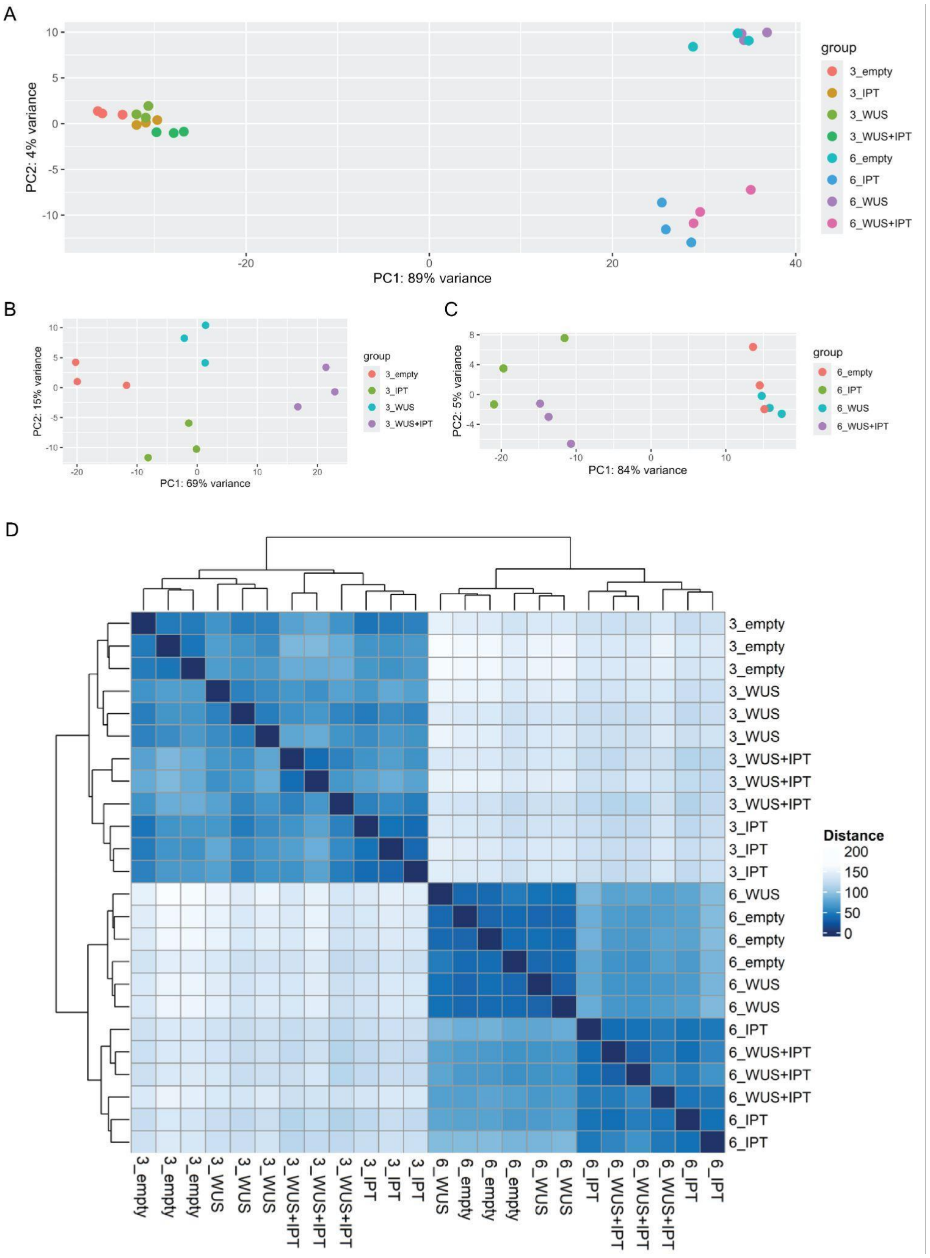


**Supplementary Figure 4:** Principal component analysis (PCA) analysis and hierarchical clustering of RNA-seq dataset. (A) PCA plots of all 3 and 6 DAT samples. (B) PCA plots of 3 DAT samples showing clustering based on treatment, whereas in 6 DAT, IPT and WUS/IPT samples were clustered together distinct from EV and WUS (C). (D) Hierarchical clustering and heat map of all of the RNA-seq samples based on gene expression similarity between samples.

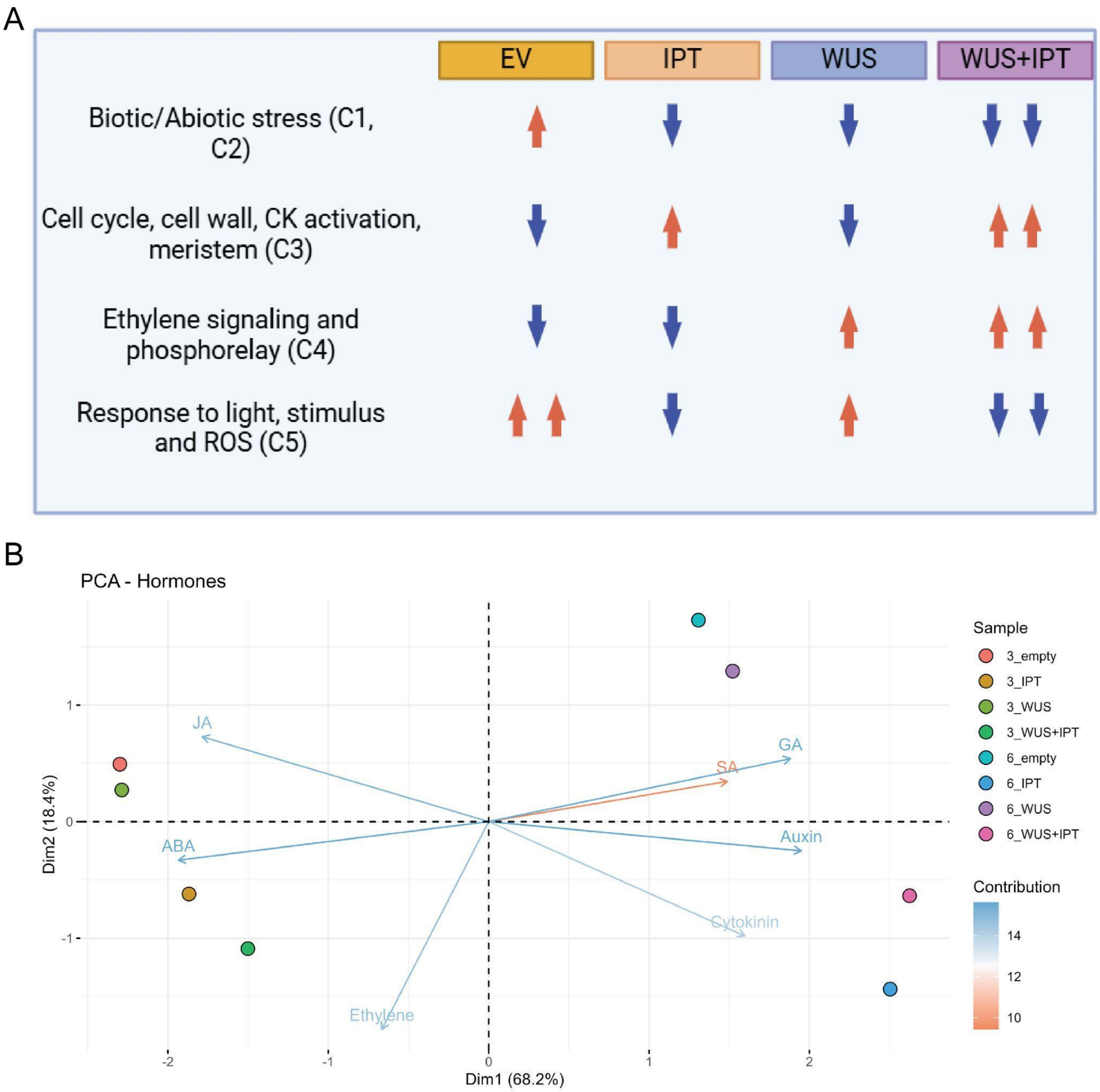


**Supplementary Figure 5:** (A) Effects of DRs on biological processes through transformation, showing upregulated (red arrows) and downregulated (blue arrows) genes. (B) Principal component analysis (PCA) plot of all samples showing genes associated with key plant growth hormones. The distribution of samples appears to be mainly based on the treatment effect. Samples distribution and scattering also suggest stronger effects of WUS and WUS/IPT on cytokinin and auxin pathways.

**Supplementary Figure 6:** Sequences of DRs cloned in T-DNA constructs.

**NosP-GmWUS-OcsT** (Red, Nos promoter. Green, *GmWUS.* Blue, OCS terminator).

gatcatgagcggagaattaagggagtcacgttatgacccccgccgatgacgcgggacaagccgttttacgtttggaactgacagaaccgc
aacgttgaaggagccactcagccgcgggtttctggagtttaatgagctaagcacatacgtcagaaaccattattgcgcgttcaaaagtcgcct
aaggtcactatcagctagcaaatatttcttgtcaaaaatgctccactgacgttccataaattcccctcggtatccaattagagtctcatattcactc
tcaatccaaataatctgcaccgtacctgcagggtccgagctaggtcacagaagcgctcaggaaggccgctgagatagaggATGATG
GAACCTCAACAACAACAACAACAAGCACAAGGGAGCCAACAACAACAACAAAACG
AGGATGGTGGCAGTGGAAAAGGGGGTTTCTGAGCAGGCAAAGTAGTAAACCACA
AAGCTCGAGAAAGGCAGAAGAAAAGGTTCACTTCTGATCATAATCATAATAATGTC
CCCATGCAAAGACCCCCAACTAATCCTTCTGCTGCTTGGAAACCTGATCTAGCTGAT
CCCATTCACACCACCAAGTATTGTAACATCTCTTCTACTGCAGGGATCTCTTCGGCAT
CATCTTCTGTTGAGATGGTTACTGTGGGACAGATGGGGAATTATGGGTATGGTTCTG
TGCCCATGGAGAAAAGTTTTAGGGACTGCTCGATATCAGCTGGGGGTAGCAGTGGC
CATGTTGGATTAATAAACCACAACTTGGGGTGGGTTGGTGTGGACCCATATAATTCC
TCAACCTATGCCAACTTCTTTGACAAAATAAGGCCAAGTGATCAAGAAACCCTTGAA
GAAGAAGCAGAGAACATTGGTGCTACTAAGATTGAAACCCTCCCTTTATTCCCTATG
CACGGTGAGGACATCCATGGCTATTGCAACCTCAAGTCTAATTCGTATAACTATGAT
GGAAACGGCTGGTATCATACTGAAGAAGGGTTCAAGAATGCTTCTCGTGCTTCCTTG
GAGCTCAGTCTCAACTCCTACACTCGCAGGTCTCCAGATTATGCTTAAGATATATCA
TATATATACTCAACTTATTAATTATCACGGTGGACTCCAACAAACGACCAGATAAGA
ATATTGAAGGAACTTTACTACAACAATGGAATTAGATCCCCGAGTGCAGAGCAGAT
TCAGAGGATCTCTGCTAGGCTGAGGCAGTACGGTAAGATTGAAGGCAAGAATGTCT
TTTATTGGTTCCAGacccagctttcttgtacaaagtggtgcctaggtgagtctagagagttaattaagacccgggactagtccct
agagtcCTGCTTTAATGAGATATGCGAGACGCCTATGATCGCATGATATTTGCTTTCAAT
TCTGTTGTGCACGTTGTAAAAAACCTGAGCATGTGTAGCTCAGATCCTTACCGCCGG
TTTCGGTTCATTCTAATGAATATATCACCCGTTACTATCGTATTTTTATGAATAATAT
TCTCCGTTCAATTTACTGATTGTACCCTACTACTTATATGTACAATATTAAAATGAAA
ACAATATATTGTGCTGAATAGGTTTATAGCGACATCTATGATAGAGCGCCACAATAA
CAAACAATTGCGTTTTATTATTACAAATCCAATTTTAAAAAAAGCGGCAGAACCGGT
CAAACCTAAAAGACTGATTACATAAATCTTATTCAAATTTCAAAAGTGCCCCAGGGG
CTAGTATCTACGACACACCGAGCGGCGAACTAATAACGCTCACTGAAGGGAACTCC
GGTTCCCCGCCGGCGCGCATGGGTGAGATTCCTTGAAGTTGAGTATTGGCCGTCCGC
TCTACCGAAAGTTACGGGCACCATTCAACCCGGTCCAGCACGGCGGCCGGGTAACC
GACTTGCTGCCCCGAGAATTATGCAGCATTTTTTTGGTGTATGTGGGCCCCAAATGA
AGTGCAGGTCAAACCTTGACAGTGACGACAAATCGTTGGGCGGGTCCAGGGCGAAT
TTTGCGACAACATGTCGAGGCTCAGCAG

**CaMV35s-IPT-OcsT** (Red, CaMV35S promoter; Green, *IPT;* Blue, OCS terminator).

tgagacttttcaacaaaggataatttcgggaaacctcctcggattccattgcccagctatctgtcacttcatcgaaaggacagtagaaaaggaa
ggtggctcctacaaatgccatcattgcgataaaggaaaggctatcattcaagatctctctgccgacagtggtcccaaagatggacccccacc
cacgaggagcatcgtggaaaaagaagacgttccaaccacgtcttcaaagcaagtggattgatgtgacatctccactgacgtaagggatgac
gcacaatcccactatccttcgcaagacccttcctctatataaggaagttcatttcatttggagaggacaATGGATCTGCGTCTAA
TTTTCGGTCCAACTTGCACAGGAAAGACGTCGACCGCGATACGTCTTGCCCAGCAGA
CTGGCCTTCCAGTCCTTTCGCTCGATCGGGTCCAATGCTGTCCTCAACTGTCAACCGG
AAGCGGACGACCAACAGTGGAAGAACTGAAAGGAACGACCCGTCTATACCTTGAAG

ATCGGCCTCTGGTGAAGGGTATCATCGCAGCCAAGCAAGCTCACGAAAGGCTGATC
GGGGAAGTGTACAATTATGAGGCCCACGGCGGGCTTATTCTTGAGGGAGGATCTAT
CTCGTTGCTCAGGTGCATGGCGCAAAGCAGTTATTGGAGTACCGATTTTCGTTGGCA
TATTATTCGCCACAAGTTAGCAGACGAGGAGACATTCATGAACGCGGCCAAGGCCA
GAGTTAGGCAGATGTTGCGCCCTGCTGTAGGCCCATCTATTATTCAAGAGTTGGTTC
ATCTTTGGAATGAGCCTCGGCTGAGGCCCATACTGAAAGAGATCGACGGATATCGA
TATGCCATGTTATTTGCTAGCCAGAACCAGATCACACCCGATATGCTATTGCAGCTT
GACCCAGATATGGAGGGTGAGTTGATTCATGGAATCGCTCAGGAGTATCTCATCCAT
GCGCGCCGGCAGGAGCAGGAATTCCCTCCAGTGAGCGTGGTCGCTTTCGAAGGATT
CGAAGGTCCACCGTTCGGAATGTGCTAGacccagctttcttgtacaaagtggtgcctaggtgagtctagagagtta
attaagacccgggactagtccctagagtcCTGCTTTAATGAGATATGCGAGACGCCTATGATCGCATG
ATATTTGCTTTCAATTCTGTTGTGCACGTTGTAAAAAACCTGAGCATGTGTAGCTCAG
ATCCTTACCGCCGGTTTCGGTTCATTCTAATGAATATATCACCCGTTACTATCGTATT
TTTATGAATAATATTCTCCGTTCAATTTACTGATTGTACCCTACTACTTATATGTACA
ATATTAAAATGAAAACAATATATTGTGCTGAATAGGTTTATAGCGACATCTATGATA
GAGCGCCACAATAACAAACAATTGCGTTTTATTATTACAAATCCAATTTTAAAAAAA
GCGGCAGAACCGGTCAAACCTAAAAGACTGATTACATAAATCTTATTCAAATTTCAA
AAGTGCCCCAGGGGCTAGTATCTACGACACACCGAGCGGCGAACTAATAACGCTCA
CTGAAGGGAACTCCGGTTCCCCGCCGGCGCGCATGGGTGAGATTCCTTGAAGTTGAG
TATTGGCCGTCCGCTCTACCGAAAGTTACGGGCACCATTCAACCCGGTCCAGCACGG
CGGCCGGGTAACCGACTTGCTGCCCCGAGAATTATGCAGCATTTTTTTGGTGTATGT
GGGCCCCAAATGAAGTGCAGGTCAAACCTTGACAGTGACGACAAATCGTTGGGCGG
GTCCAGGGCGAATTTTGCGACAACATGTCGAGGCTCAGCAG

**CaMV35s-GmGRF4-GmGIF1-OcsT** (Red, CaMV35S promoter; purple, GmGRF4; green, GIF; Blue, OCS terminator).
tgagacttttcaacaaaggataatttcgggaaacctcctcggattccattgcccagctatctgtcacttcatcgaaaggacagtagaaaaggaa
ggtggctcctacaaatgccatcattgcgataaaggaaaggctatcattcaagatctctctgccgacagtggtcccaaagatggacccccacc
cacgaggagcatcgtggaaaaagaagacgttccaaccacgtcttcaaagcaagtggattgatgtgacatctccactgacgtaagggatgac
gcacaatcccactatccttcgcaagacccttcctctatataaggaagttcatttcatttggagaggacaATGAACATCAGTGGC
GGAGGAGGAACTGTGATGGGTTTCAGTAGTAATGGGAGGTCCCCATTCACAGTGTCT
CAGTGGCAGGAACTGGAGCACCAAGCTTTGATTTTCAAGTACATGGTTGCGGGTCTT
CCTGTGCCTCCCGATCTCGTCCTCCCCATTCAGAAGAGCTTCGACTCTACTCTCTCTC
ACGCTTTCTTTCACCATCCCACACTGAGTTATTGTTCCTTCTATGGGAAGAAGGTGGA
CCCTGAGCCAGGACGATGCAGGAGGACTGATGGAAAAAGTGGAGGTGCTCCAAG
GAAGCATACCCAGACTCCAAGTACTGCGAGCGCCACATGCACCGTGGCCGCAACCG
TTCAAGAAAGCCTGTGGAATCACAAACTATGACTCACTCGTCTTCAACTGTCACATC
ACTCACTGTGACTGGTGGTGGTGACAGTAATGGAACTGTAAACTTCCAAAACCTTCC
CACAAATGCCTTTGGTAATCTCCAGGGTACTGATTCTGGAACTGACCGCACGAATTA
TCATCTAGATTCCATTCCCTATGCGATTCCAAGTAAAGAATACAGGTGTCTTCAAGG
ACTTAAATCTGAGGGTGGTGAACACTGCTTCTTTTCTGAAGCTTCTGGAAGCAACAA
GGTTCTCCAAATGGAGTCACAGCTGGAAAACACATGGCCTTCGATGTCAACCAGAG
TTGCCTCTTTTTCTACATCAAAATCAAGTACTGATTCCCTGTTGCATAGTGATTATCC
CCAACATTCGTTTTTATCTGGTGAATATGCATCGGGAGAACACGTGAAGGAGGAGG
GCCAGCCTCTTCGACCTTTTTCTAATGAATGGCCTAAAAGCAGGGAGTCATGGTCTG
GTCTGGAAGATGATATATCCAACCAAACAGCCTTCTCCACAACTCAACTCTCAATAT

CCATTCCTATGTCTTCCGATTTCTCTGCAACGAGCTCTCAGTCCCCACATGGTGAGAA
TGAGATTCAATTTAGG<u>GCGGCCGCTGCC</u>ATGCAGCAGCACCTGATGCAGATGCAGCCC
ATGATGGCTGCCTACTACCCCAACAACGTCACCACTGATCACATTCAACAGTACCTG
GATGAGAACAAGTCCTTGATTCTGAAGATTGTTGAAAGCCAGAATTCTGGCAAGCTG
AGCGAGTGTGCCGAGAACCAATCAAGGCTGCAGAGAAATCTCATGTACCTAGCTGC
AATAGCTGATTCTCAACCACAACCATCTCCATTGGCTGGTCAGTATCCTTCTAGTGG
ACTTGTGCAGCAAGGAGCACACTACATGCAGGCTCAACAGGCTCAGCAGATGTCAC
AACAACAGCTAATGGCTTCGCGCTCCTCGCTCCTGTACTCCCAACAGCCTTTCTCAG
TGCTTCAACAGCAGCAAGGCATGCACAGCCAACTTGGCATGAGCTCCAGTGGAAGT
CAAGGCCTCCACATGCTGCAAAGTGAAGCCACTAATGTTGGAGGCAATGCAACCAT
AGGAACCGGAGGAGGGTTTCCGGACTTTGTACGCATTGGTAGTGGCAAGCAAGATA
TTGGAATCTCTGGTGAAGGCAGAGGAGGAAACTCTAGTGGCCACTCTGGTGATGGT
GGTGAGACACTTAATTACCTGAAAGCTGCTGGTGATGGAAACTGAacccagctttcttgtacaaa
gtggtgcctaggtgagtctagagagttaattaagacccgggactagtccctagagtcCTGCTTTAATGAGATATGCGAG
ACGCCTATGATCGCATGATATTTGCTTTCAATTCTGTTGTGCACGTTGTAAAAAACCT
GAGCATGTGTAGCTCAGATCCTTACCGCCGGTTTCGGTTCATTCTAATGAATATATC
ACCCGTTACTATCGTATTTTTATGAATAATATTCTCCGTTCAATTTACTGATTGTACC
CTACTACTTATATGTACAATATTAAAATGAAAACAATATATTGTGCTGAATAGGTTT
ATAGCGACATCTATGATAGAGCGCCACAATAACAAACAATTGCGTTTTATTATTACA
AATCCAATTTTAAAAAAAGCGGCAGAACCGGTCAAACCTAAAAGACTGATTACATA
AATCTTATTCAAATTTCAAAAGTGCCCCAGGGGCTAGTATCTACGACACACCGAGCG
GCGAACTAATAACGCTCACTGAAGGGAACTCCGGTTCCCCGCCGGCGCGCATGGGT
GAGATTCCTTGAAGTTGAGTATTGGCCGTCCGCTCTACCGAAAGTTACGGCACCAT
TCAACCCGGTCCAGCACGGCGGCCGGGTAACCGACTTGCTGCCCCGAGAATTATGC
AGCATTTTTTTGGTGTATGTGGGCCCCAAATGAAGTGCAGGTCAAACCTTGACAGTG
ACGACAAATCGTTGGGCGGGTCCAGGGCGAATTTTGCGACAACATGTCGAGGCTCA
GCAG
